# High-resolution label-free transcranial imaging of *in vivo* neural activity via interferometric measurement of tissue deformation

**DOI:** 10.1101/2023.10.05.561052

**Authors:** Austen T. Lefebvre, Carissa L. Rodriguez, Eyal Bar-Kochba, Nicole E. Steiner, Marek Mirski, David W. Blodgett

## Abstract

Rapid sub-nanometer neuronal deformations have been shown to occur as a consequence of action potentials *in vitro*, allowing for registration of discrete axonal and synaptic depolarizations and thus providing a novel signature for recording neural activity (1–3). We demonstrate that this signature can be extended to *in vivo* measurements through recording of rapid neuronal deformations on the population level with optical phase-based recordings. Complicating these measurements is the optical phase noise due to microvascular flow as well as the presence of significant tissue clutter (deformation) associated with physiologic processes (e.g., heart and respiratory rate). These recordings were acquired using a full-field holographic imaging system with spatiotemporal resolutions of less than 1 ms and 0.1 mm^3^ over a 3 mm diameter field of view (FOV). Our system demonstrates, for the first time, the ability to non-invasively record *in vivo* tissue deformation associated with population level neuronal activity. We confirmed this technique across a range of neural activation models, including direct epidural focal electrical stimulation (FES), activation of primary somatosensory cortex via whisker barrel stimulation, and pharmacologically-induced seizures. Calibrated displacement measurements of the associated tissue deformations provided additional insight into the underlying neural activation mechanisms. Collectively, we show that holographic imaging provides a pathway for high-resolution, label-free, non-invasive recording of transcranial *in vivo* neural activity at depth, making it highly advantageous for studying neural function and signaling.

## Introduction

High-resolution, non-invasive recording of neural activity has been a goal of clinical and investigative neurosciences to better understand human cognitive function and to improve neuro-therapeutics (4). Investigations have invoked numerous technologies to non-invasively record and map neuronal signaling (4, 5), yet lacking is a technology having both spatial and temporal resolutions to match the discrete phenomena of axonal and synaptic depolarizations evaluated in a dynamic, *in vivo* model. Electrical recordings have long been the gold standard and are capable of invasively recording activity from the single neuron level up to population level via electrodes (i.e. electrocorticography, electrode pair and/or patch clamp techniques) (6). Non-contact, label-free methods, however, are advantageous to avoid damage to the neural tissue as well as to enable greater flexibility in recording neural activity.

For decades, optical recordings have successfully detected changes that accompany neuronal activation (7, 8), suggesting a photonics approach may be feasible for non-contact, label-free neural activity detection and imaging. Numerous studies have investigated intrinsic optical changes coincident with neural activation, including alterations in tissue reflectivity or intensity, and polarization changes (9). Another promising approach to realize this goal is to directly measure neuronal deformations occurring during the action potential. Researchers have recorded full-field, spike-induced deformations in electrogenic cells through spike averaging (10–13). This work demonstrated that successful recording of these deformations requires a high spatial resolution imaging system with sub-nanometer displacement sensitivity at sub-millisecond temporal resolution (12, 13). Recent studies aimed at detecting these displacement signals utilize coherent optics-based approaches because of the high sensitivity afforded by phase-based measurements to provide insight into tissue dynamics (e.g., deformation) (14). Phase-based measurements have been successfully *demonstrated in vitro* with plated cells as well as *ex vivo* with dissected squid giant axon (10–13, 15–17). These results show that tissue dynamics and electrical recordings of neural activity are well correlated.

Extending these optical phase-based approaches to *in vivo* models is challenging, primarily due to the increased phase noise incurred by the intrinsic temporal dynamics of living tissue. The physiological artifacts derived from respiration and blood flow, for example, are considerable. Phase noise limits the measurement sensitivity, impacting the ability to detect the nanometer-scale displacements associated with neuronal activity. One of the dominant sources of phase noise for *in vivo* measurements is the decorrelation time of the tissue, which is attributed to correlated (e.g., respiratory) and uncorrelated (e.g., microvascular) motion of the scatterers in the tissue itself. The decorrelation time for *in vitro* samples is on the order of seconds whereas for *in vivo* samples is on the order of milliseconds (18–20), thus greatly increasing recording challenges.

Our research has focused on the development of a full-field digital holographic imaging (DHI) system that minimizes optical phase noise to enable *in vivo*, transcranial non-invasive detection of population level cortical activity. Three mechanistically distinct neural activation models were used to validate this technique: direct epidural focal electrical stimulation (FES), intrinsic activation of primary somatosensory cortex via whisker deflection, and pharmacologically-induced cortical seizure activity. This is the first demonstration of such optical phase detection of cortical activity performed *in vivo*, illustrating the spatiotemporal progression of population-level neural activation through both cranial window preparations and at-depth through intact cranium.

### Digital Holographic Imaging for *in vivo* neural activity recording

The DHI system presented here measures, with a high degree of sensitivity, the sub-nanometer, out-of-plane deformation of tissue sample over time, providing a means to measure population level neuronal activation. The optical system consists of an interferometer that illuminates the cortical region with a coherent light source (Fig. 1A). The scattered light from the cortex is coherently mixed with a reference beam, which is derived from the same light source, at the focal plane array (FPA) in an off-axis holography configuration. The coherent mixing forms an interference pattern that is referred to as a hologram, which can be digitally processed to compute the tissue deformation. See Materials and Methods for more details about the DHI system.

**Fig 1.**
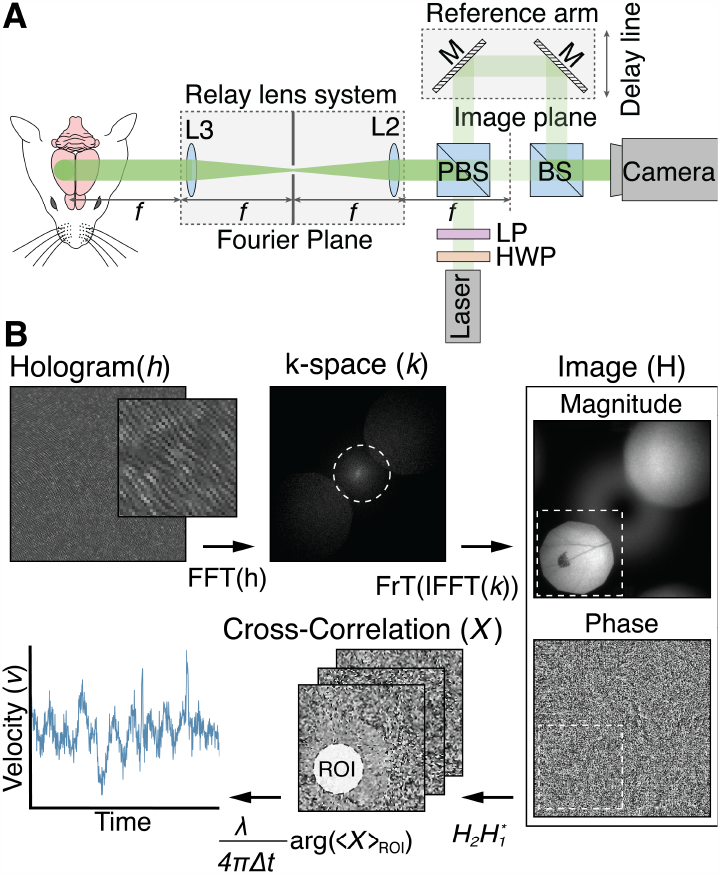
Overview of digital holographic imaging (DHI) system, image reconstruction and velocity calculation. (A) DHI system schematic. L = lens, PBS = polarizing beam splitter, BS = beam splitter, M = mirror, HWP = half wave plate, LP = linear polarizer. A collimated laser is incident on a PBS. The reflected beam illuminates the sample forms an image behind the PBS through a lens system. The reference beam transmits directly through the PBS, into a variable-length delay line and combines with the object light via a 10/90 BS. A camera FPA records a hologram. (B) Reconstruction overview to compute target velocity. A Fourier transform (FFT) is applied to the hologram for conversion to k-space and filtered at a cutoff frequency (dashed line) to mitigate the DC term. The complex image with magnitude and phase is reconstructed by applying an inverse FT (IFFT) and Fresnel transform (FrT) and subsequently cropped to an area containing the real image (dashed line). A cross-correlation image between time subsequent time points is computed without a spatial shift and a processing region-of-interest (ROI) is defined. The average velocity within the ROI is computed by spatially averaging complex pixels in the ROI and scaling by wavelength (*λ*) and sampling interval (Δ*t*) to convert from radians to units of velocity.

Computing the tissue deformation is a multistep process that starts with filtering the DC term from the hologram and subsequently numerically propagating the wavefront to the depth of the cortex using a Fresnel transform. (Fig. 1B). The resulting image (*H*) contains magnitude and phase information of the sensing volume captured at rates greater than 1.5 kHz, which helps minimize optical phase noise due to the millisecond decorrelation time of the cortical tissue (19). Phase change is computed by spatial averaging within a user-defined region-of-interest (ROI) to reduce phase error due to uncorrelated phase noise among pixels within the ROI, allowing us to resolve nanometer scale frame-to-frame displacements. The spatial averaging procedure is performed by first computing a cross-correlation between two subsequent images, given by:

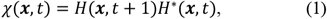

where ***x*** and *t* denote the spatial and time dimensions, respectively, and * is the complex conjugate operation. The average phase change within an ROI is given by:

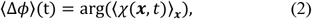

where ⟨.⟩_***x***_ is the independently averaged real and imaginary components of χ. The tissue velocity is then computed by scaling ⟨Δ*ϕ*⟩ by the wavelength (*λ*) of the light and sampling interval (Δ*t*), given by:

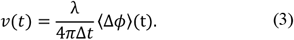

The corresponding displacement was computed by numerically integrating the velocity. The spatial averaging defined in the complex domain has two benefits. First, the phase change on a per-pixel basis is weighted by the magnitude product of the two images, thus down weighting phase values with high noise due to low magnitude values. Secondly, averaging in the complex domain does not suffer from 2π wrapping issues, in contrast to averaging phase in the real domain (21).

Two different DHI systems were fabricated and designed to operate at 780 nm and 1310 nm. Operation at a shorter wavelength provides better velocity sensitivity, so the 780 nm system was selected for baseline recordings to verify that the system can achieve the required performance. The second 1310 nm system achieves improved at-depth sensing performance by reducing photon propagation loss due to optical scatter (22), as required for the transcranial measurements. Both systems were designed to provide an intrinsic spatial resolution of approximately 30 μm. Depth resolution is provided through the use of a short coherence length laser. The 780 nm system utilized a 3 mm coherence length laser and the 1310 nm system utilized a 50 μm coherence length laser, which provided depth resolutions of approximately 1 mm and 16 μm, respectively. The location of this coherence gate is controlled by a delay line in the reference arm, with the goal of placing this sensing volume within the region of measured neural activity to maximize recording sensitivity.

Validation of this methodology was demonstrated through a variety of experiments designed to precisely measure the temporal and spatial characteristics of the tissue deformation associated with neuronal activation, and to demonstrate the high sensitivity and specificity of *in vivo* optical recordings of cortical neural activation. The first set of experiments explored FES for intact dura mater recordings, due to its well-characterized spatiotemporal response. Successful single-shot recordings of FES epidural stimulation motivated additional transcranial measurements to demonstrate at-depth capabilities of the approach. Additionally, we investigated the ability to detect intrinsically evoked responses in the primary somatosensory barrel cortex (wS1) via thalamo-cortical efferents triggered by mechanical whisker deflection. Lastly, we made measurements of tissue deformation during cortical seizure discharges to characterize the larger spatial extent and synchronous neural activation associated with ictal neural activity. Results of the tissue deformations are presented from single stimulations and with stimulus-locked averaging (SLA) to reduce the phase noise, as well as with corresponding electrophysiological measurements.

## Results

### Tissue deformation associated with neural activity is modulated by FES current

Initial FES measurements were made with the 780 nm DHI system with a 3 mm coherence length laser. FES was exacted using a pair of tungsten stimulating electrodes placed on the exposed epidural surface (Fig. 2A and B) following hemicraniectomy of anesthetized, paralyzed, and ventilated Sprague-Dawley rats (350 – 450 g). Epidural (versus subdural) stimulation was utilized to minimize trauma to cortical tissue. Repetitive cathodic bipolar stimulations presented at 10 Hz were performed via a constant current isolated pulse stimulator at currents of 0.25 mA, 0.50 mA, and 0.75 mA, and neural responses were recorded via DHI.

**Fig. 2.**
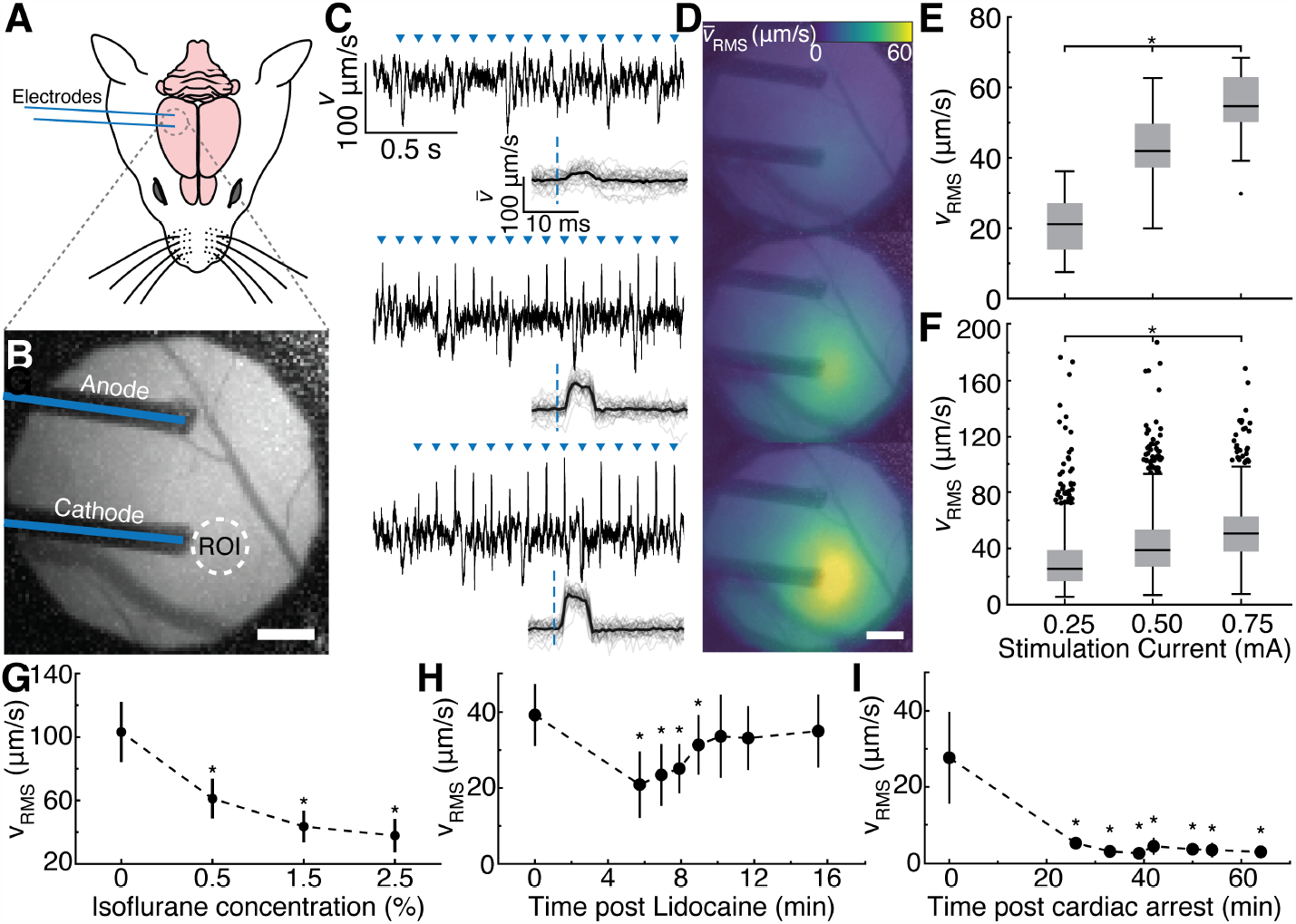
Cortical tissue velocity following focal epidural electrical stimulation (FES). (A) Rodent schematic setup for FES showing electrodes placed on top of dura mater. (B) Average magnitude image of the cortex with electrodes (blue lines) and ROI (white dotted circle) identifies the pixels averaged for the velocity waveforms. Scale bar, 0.5 mm. (C) Velocity waveforms at (top) 0.25 mA, (middle) 0.50 mA, and (bottom) 0.75 mA FES current with FES onset indicated by blue triangles. Insets show the velocity with stimulus-locked averaging (SLA) (*n* = 30 events; black line). (D) Maps of the Root mean square (RMS) velocity with SLA (*n* = 30 events) within 10 ms following FES for (top) 0.25 mA, (middle) 0.50 mA, and (bottom) 0.75 mA FES current, computed using a 0.5 mm diameter sliding window. Average magnitude image is overlaid with transparency on the velocity map. (E) RMS velocity within 10 ms following FES for rodent shown in C and D (*n* = 30). (F) RMS velocity for multiple rodents (0.25 mA: *n* = 617 stimulations for 6 rodents; 0.50 mA: *n* = 1,118 stimulations for 11 rodents; 0.75 mA: *n* = 1,160 stimulations for 7 rodents). Box plots indicate the median (black line), 25% and 75% quantiles (gray box), non-outlier extrema (whiskers), and outliers defined as 1.5 times the interquartile range (black circle). *p < 0.05 (Kruskal-Wallis with post-hoc Dunn–Šidák correction for multiple comparisons). RMS velocity within 10 ms following administration of (G) Isoflurane and (H) Lidocaine, and (I) induction of cardiac arrest. Results are shown as mean (black circle) ± standard deviation from *n* = 30 FES events. *p < 0.05 (one-way ANOVA with post hoc Dunn–Šidák correction for multiple comparisons).

Velocity waveforms extracted from an 0.5 mm ROI adjacent to the cathode reveal the onset of the optically detected changes in tissue velocity associated with evoked neural activation within 5 ms of stimulation onset (Fig. 2C). At higher currents of 0.50 mA and 0.75 mA, the response was visible at a single stimulation level, whereas at 0.25 mA SLA (*n* = 30 events) was needed to reduce the velocity noise. The spatial distribution of the tissue velocity as quantified by the root mean square (RMS) of the velocity with SLA (*n* = 30 events) within 10 ms following FES shows that the activation area increases with greater current (Fig. 2D). The corresponding activation magnitude, quantified as the RMS velocity within 10 ms following stimulus within a 0.5 mm ROI adjacent to the cathode, significantly increased with stimulation current (*p* < 0.05; Fig. 2E and F), with the majority of RMS velocities falling within the range of 5 µm/s – 100 µm/s. Comparing RMS velocity from a single rodent (Fig. 2E) to multiple rodents (Fig. 2F), there was more overlap in RMS velocity between stimulus currents.

The source of the detected changes in tissue velocity associated with neuronal activation was demonstrated, via specifically targeted control experiments with pharmacologic synaptic inhibition. In the first model, direct cortical neuronal inhibition was implemented via induction of general anesthesia using isoflurane. Isoflurane anesthesia resulted in a statistically significant dose-dependent diminution of the post-stimulus RMS velocity compared to the pre-anesthetic baseline condition of 0% isoflurane concentration (*p* < 0.05; Fig. 2G). In the second anesthetic model, topically applied lidocaine (4%), a sodium channel blocker, temporarily reduced the ability for FES to yield evoked changes in tissue velocity as shown by the statistically significant decreases in the RMS velocity compared to the pre-anesthetic baseline (*p* < 0.05; Fig. 2H). Lastly, a post-cardiac arrest model was conducted to assess the tissue velocity in the absence of viable cellular function. The RMS velocity significantly decreased relative to the baseline condition (*p* < 0.05) following circulatory failure (Fig. 2I).

### DHI sensitivity to FES induced neuronal deformations is maintained through intact cranium

An FES model of cortical activation was used to evaluate the system’s sensitivity to measure transcranial tissue velocities. For this model, the electrodes were placed in contact with the dura mater through 2 mm burr holes made through the intact cranium (Fig. 3A and B). Tissue velocities were evaluated within a 0.5 mm ROI that did not overlap with the burr holes. Pre-stimulus Z-score normalization was applied to the segmented velocity prior to SLA (*n* = 30 events) to account for the effects of optical scattering while imaging at depth. For these measurements, the 1310 nm DHI system was selected using a 50 µm coherence length laser, to improve at-depth sensing and depth resolution, respectively.

**Fig. 3.**
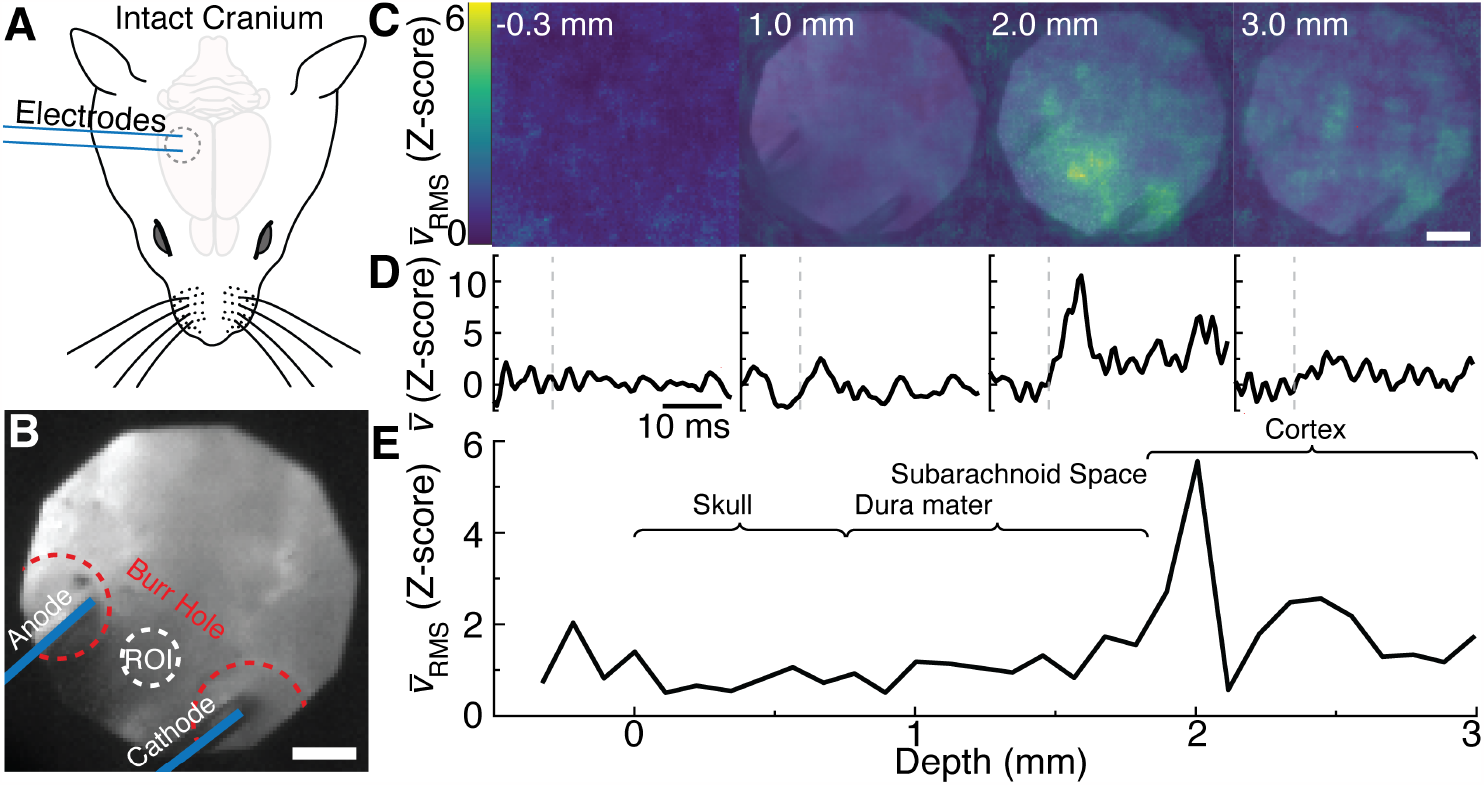
Transcranial cortical tissue velocity following FES. (A) Schematic of the rodent setup showing through-cranium electrode placement for FES at 0.75 mA. (B) Average magnitude image of the cortex showing the placement of electrodes (indicated by blue lines) in contact with the dura mater through burr holes made in the cranium (perimeter indicated by red dotted circles). The ROI (white dotted circle) identifies the pixels averaged together to obtain the velocity waveforms for (D) and (E). (C) Maps of the RMS velocity with SLA (*n* = 30 events) with pre-stimulus Z-score normalization following FES for imaging depth of -0.3 mm to 3 mm (left to right). Maps were computed using a 0.5 mm diameter sliding window. The average magnitude image is overlaid with transparency on the velocity map. Scale bar, 0.5 mm. (D) Pre-stimulus Z-score normalized velocities with SLA (*n* = 30 events) with FES onset denoted as a vertical dashed line for depth shown in (E). (E) RMS velocity with SLA (*n* = 30) with pre-stimulus Z-score normalization for ROI shown in (B) as a function of depth with the approximate ranges of depth for the anatomical layers shown. Reported depths are relative to the surface of the cranium.

Maps of the RMS velocity demonstrated the system’s ability to measure and localize neural activity at depths approximately at the surface of the cortex (i.e., 2.0 mm) (Fig. 3C), where the RMS velocity within 10 ms is approximately 6 standard deviations above the pre-stimulus mean between the electrodes. The temporal response of the velocity with SLA at depths of 2.0 mm shows a less than 5 ms onset post-stimulus with durations of 10 ms (Fig. 3D), similar to the optical response measured in the exposed cortex (Fig. 2B inset). In contrast, at shallower depths within the cranium, dura mater, and subarachnoid space, the RMS velocity response is less than 2 standard deviations above the pre-stimulus baseline (Fig. 3E). These different depth recordings were achieved by varying the relative location of the coherence gate through control of the optical delay line in the DHI system’s reference arm.

### Intrinsic cortical stimulation elicits a neural deformation response

Optical recording of neural activity was extended to *in vivo* models of intrinsic evoked cortical activity to demonstrate the ability of this technique to extend to intrinsic cortical stimulation paradigms. In the first model of evoked activity, we measured tissue velocity in the primary somatosensory barrel cortex (wS1) following single whisker deflection (Fig. 4A). As with the transcranial FES recordings, these measurements were acquired using the 1310 nm optical system with a laser coherence length of 3 mm (instead of 50 µm for transcranial measurements). Imaging of wS1 through a transparent electrocorticography (ECoG) grid was employed to provide a simultaneous temporal comparison of neural activity (Fig. 4B). A high-pass filter was applied to the velocity data to remove lower frequencies that occur from physiologic-related artifacts, e.g., due to blood flow or respiration. Similarly, high-gamma bands were computed from the ECoG data, as to compare similar frequency content between the optical and electrical data. SLA (*n* = 50 events) of ECoG and tissue velocity demonstrated nearly coincident temporal onset of approximately 10 ms and duration of approximately 50 ms (Fig. 4C and D). Comparing across 5 rodents, we found statistically significant differences (*p* < 0.05) in the pre-to post-stimulus RMS velocity with SLA (*n* = 30 events) (Fig. 4E). These differences between the pre-to-post stimulation velocities were still significant (*p* < 0.05) for all but a single rodent without SLA (Fig. 4F). The range of velocities in the wS1 ranged between approximately 0.2 µm/s – 8 µm/s.

**Fig. 4.**
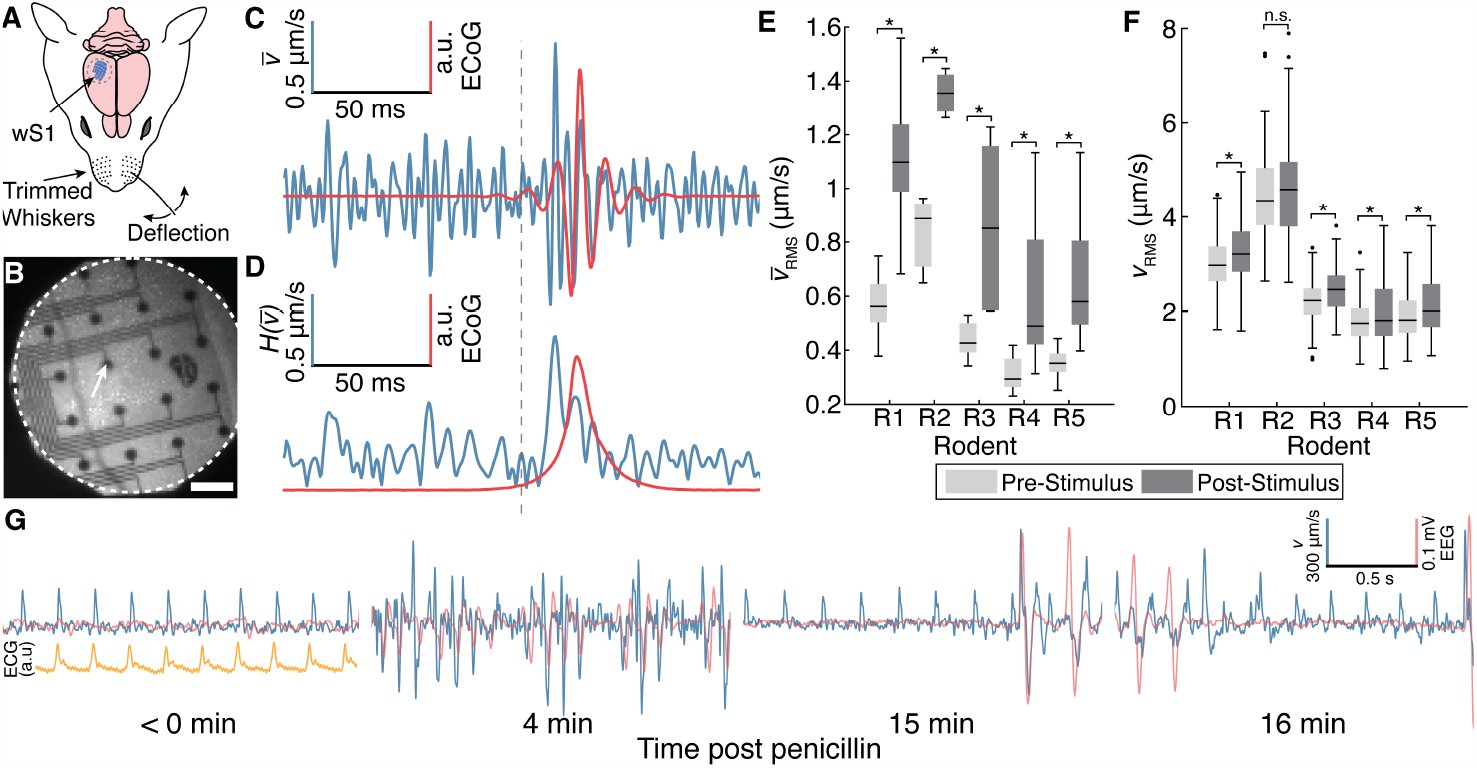
Cortical tissue velocity associated with intrinsically evoked neural activity. (A) Rodent setup for single whisker deflection and primary barrel cortex (wS1) imaging. (B) Magnitude image of wS1 with epidural ECoG grid. Scalebar, 0.5 mm. (C) Velocity with SLA and corresponding (D) Hilbert envelope (n = 50 stimulations; blue) overlayed with ECoG waveforms (n = 50 stimulations; red) from electrode indicated by white arrow in (B). (E) RMS velocity with SLA for 5 rodents (R1: *n* = 9; R2: *n* = 3; R3: *n* = 7; R4: *n* = 21; R5: *n* = 13) computed 25 ms pre-stimulus (light gray) and 25 ms post-stimulus (dark gray). Each data in the box is computed from 30 stimulus-locked events. (F) Corresponding pre- and post-stimulus RMS velocity without event averaging for 5 rodents (R1: *n* = 296; R2: *n* = 99; R3: *n* = 210; R4: *n* = 634; R5: *n* = 399) .*p < 0.05 (Mann-Whitney U test). (G) Velocity (blue) and EEG (red) waveforms for a rodent before and 4 min, 15 min, and 16 min after application of topical penicillin on rodent dura mater to induce seizure. Inset in (G) shows the associated electrocardiogram (ECG; orange) waveform.

The second method of naturally evoked neural activity was achieved with topical penicillin, a well–known GABA antagonist and convulsant model of spike-wave epilepsy (24). These measurements were acquired using the 780 nm optical system. Following topical application of penicillin to the cortex, spontaneous and repetitive electroencephalogram (EEG) sharp waves/spikes ranging between 75 µV – 250 µV were observed synchronously with tissue velocity that ranged between ±500 µm/s (Fig. 4G). The agreement with the velocity measurement was especially pronounced in concert with high-voltage ballistic EEG sharp waves.

### Tissue deformations depend on neural stimulation model

To compare tissue deformations across the neural stimulation models, we computed the total tissue displacement post-stimulation. We found that tissue displacements ranged from a median of 6.7 nm with wS1 activation to 5.9 μm with ictal discharges (Fig. 5). The median displacements associated with FES for stimulation currents of 0.25 mA, 0.5 mA, and 0.75 mA were 91.2 nm, 140.9 nm, and 190.2 nm respectively. Across these models, we found significant differences (*p* < 0.05).

**Fig. 5.**
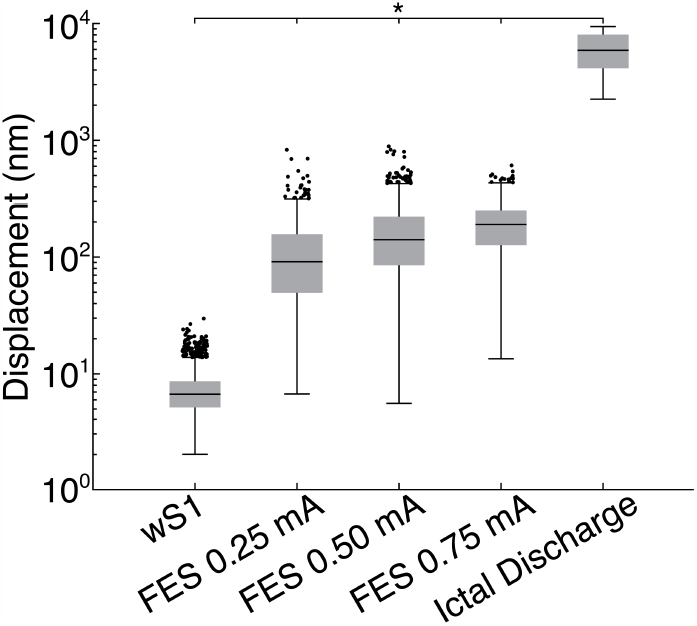
Total tissue displacement across a range of neural stimulation paradigms. wS1: *n* = 1638; FES 0.25 mA: *n* = 617; FES 0.50 mA: *n* = 1,118; FES 0.75 mA: *n* = 1,160; Ictal Discharge: *n* = 20. *p < 0.05 (Kruskal-Wallis with post-hoc Dunn–Šidák correction for multiple comparisons).

## Discussion

The results show the ability of DHI to effectively resolve changes in tissue deformation associated with neural activity, which were validated through a variety of simulation paradigms designed to precisely measure the spatiotemporal characteristics of the deformations.

Initially, measurements were made by stimulating cortical neurons using epidural FES through an intact dura, as it is the most direct model of neural activation. This FES approach ensured a repeatable and synchronized activation of neuron populations. We observed neural tissue deformation (as measured by velocity) on a single stimulation level (i.e., single-shot) for the 0.5 mA and 0.75 mA stimulation current, demonstrating, that such measured deformations are distinct, and separable from physiologic tissue deformation due to non-synaptic physiologic processes or intrinsic noise of the DHI system. Decreasing the stimulation current to 0.25 mA required SLA to diminish the contribution of these noise sources, and thereby effectively register neural activity-derived cortical tissue deformation. Spatial velocity maps (Fig. 2D) localize the tissue deformation proximate to the cathode, supporting the concept of the preferential activation of axons (25, 26) rather than somas or terminals, as is more commonly observed with anodic stimulation (27). The spatial extent of tissue deformation varied from 0.1 mm to 0.5 mm for stimulation currents of 0.25 mA to 0.75 mA, respectively, and propagated radially from the cathode within approximately 2 – 3 ms (Fig. 2D). These activation areas and propagation times of tissue deformation are consistent with estimates of current spread *in vivo* (28–30), suggesting that this measurement is from an aggregated neuromechanical response of a population of neurons and that the recruitment of the neurons increases with stimulation current (Fig. 2D and E). However, capturing an accurate estimate of neuronal recruitment is challenging, given that the DHI system measures the average tissue deformation within the sensing depth, which was approximately 1 mm, based on the coherence length of the light source. Assuming radial symmetry of the neural activation, this sensing length is greater than the activation length as defined by the height of the activated neuronal layer, resulting in a blurred response. To minimize this blurring effect, the sensing volume should be smaller than the activity volume, which would be achieved by further reducing the coherence gate.

Cortical tissue motion following epidural FES is also impacted by the underlying neurophysiologic state. We found reductions in tissue velocity following direct cortical neuronal inhibition with both systemic (isoflurane) and local (lidocaine, 4%) anesthesia (Fig. 2G and H). These results demonstrate that retarding the depolarization of cortical neurons decreases aggregate neuronal tissue motion. The dose-dependent decline in tissue velocity with increasing concentrations of isoflurane further demonstrates tissue motion as a measure of the number of neurons depolarizing. In addition, the approximate 10-minute decrement in response following topical lidocaine, offers excellent alignment with published pharmaco-dynamic inhibition via iontophoretically-applied lidocaine to cortical neurons (31). Utilizing a post-cardiac arrest model, we further observed a decrement in velocity over time as the number of viable neurons gradually diminished as corroborated by EEG (Fig. 2I). These results further highlight the direct relationship of our recorded tissue deformation to quantitative neurons depolarization. Understanding other factors that are likely involved should be answered with future research. As an example, we suspect that synchronicity of neuronal depolarization is an important factor, but we are unable to make definite conclusions based on the presented data.

Beyond imaging the superficial cortical response during epidural FES, we explored measuring neural activity at moderate depths. Utilizing epidural FES, we measured cortical tissue deformation through the intact rat cranium at high spatiotemporal resolutions (Fig. 3). Greater depth penetration was achieved by increasing the wavelength to 1310 nm, thus reducing optical scatter (32), while greater depth resolution of approximately 16 μm was achieved by decreasing the coherence length of the laser to 50 µm. By progressively imaging through depth, we found that neural tissue deformation was maximized near the estimated surface of the cortex at a depth of approximately 2 mm below the surface of the cranium. Unlike the results for FES with intact dura, the maximum tissue deformation was concentrated between the anode and the cathode. This discrepancy is attributed to the difficulty in precisely controlling the orientation of the DHI system relative to the cortex surface, resulting in a sensing volume that is not parallel to the cortex surface. Furthermore, since epidural FES preferentially activates neurons near the surface of the cortex, it limited our ability to characterize DHI system performance through greater neural tissue depths. Future studies utilizing depth electrodes for FES will be needed to assess the penetration capabilities of DHI. Despite these limitations, these results demonstrate the feasibility of DHI for non-invasive, label-free detection of neural activity through scattering tissue, which overcomes the invasiveness required for measuring *in vivo* neural activity with approaches such as electrophysiology or fluorescence imaging.

To expand the utility of DHI for measuring *in vivo* neural activity, we moved from direct activation with FES to intrinsically evoked cortical activation due to whisker deflection and seizures. By imaging wS1 following single whisker deflection, we found that the tissue deformation had an onset and duration similar to simultaneously collected ECoG recordings (Fig. 4C). However, tissue velocities were substantially reduced than velocities induced by FES. To effectively extract these reduced tissue velocities, we applied a high-pass filter with a cutoff frequency of 100 Hz to suppress the physiological artifacts, and leveraged SLA to reduce both physiological and system phase noise. It is important to note that the SLA approach also reduced the measured post-stimulus velocity. However, this reduction was to a lesser degree then the pre-stimulus velocity, indicating that the post-stimulus velocities had more variability than those the induced velocities due to FES. We attribute the differences in velocity between FES and whisker stimulation to the unique structural organization and function of the wS1. Unlike FES, where direct and synchronous neuronal activation occurs, the neural activity due to whisker stimulation propagates to wS1 via primary thalamic connectivity, then via thalamo-cortical pathways. Since the sensing depth was approximately 1 mm, the tissue deformation linked to the neural propagation through these layers is most likely blurred. To more accurately map the tissue deformation in wS1, additional studies are needed with a shorter coherence length laser that progressively image layers of wS1. Despite these limitations, we were able to observe significant differences between pre- and post-stimulus velocities across a cohort of multiple rodents (Fig. 4E and F). To complement the wS1 model with another specific mode of neural activation, we used a seizure model to study the tissue deformation associated with an intrinsically evoked activation source that has a larger spatial extent and synchronous activation (33). We found that the tissue deformation during ictal discharges was similar to measurements made with EEG (Fig. 4G), and that these velocities were substantially higher than those induced by FES and whisker stimulation. We attribute these high velocities to synchronous recruitment of a large quantity of neurons within the sensing volume.

Collectively, our findings show that cortical tissue deformation associated with *in vivo* neural activity is dependent on the mode and intensity of neural activation. Although in this study we examined tissue deformation using velocity, for comparison to previous *in vitro* and *ex vivo* studies (1, 2, 34), we also computed the total displacement of tissue during stimulation events (Fig. 5). Consistent with trends observed with tissue velocity, the smallest displacements occurred during whisker stimulation, followed by FES (scaled by current), and the largest displacements occurred during ictal discharges. Compared to previous *in vitro* or *ex vivo* studies, where the neuronal displacements are on the order of a few nanometers, the measured *in vivo* displacements are substantially larger. These differences are perhaps expected, given that the aggregate population response of neurons will be influenced by a variety of factors, such as the total quantity, density, and orientation of neurons, as well as the activation synchronicity. Although there has been significant progress in modeling the coupled electromechanical response of single neurons and nerves, as reviewed by Peets et al. (35), scaling these models to neural populations is critical to understand the physical underpinnings that bring rise to the bulk tissue deformations associated with neural activation.

These findings open new avenues for optical recording of brain function by establishing DHI measurements of neural deformation as a label-free modality with high spatiotemporal resolution capable of resolving neural activity complementary to existing techniques with strong potential for both basic and clinical neuroscience applications.

## Materials and Methods

### Animal models and preparation

Sprague-Dawley male rats (325 – 450 g; Charles River, Wilmington, MA) were utilized for *in vivo* experiments. All experimental procedures were performed in accordance with approved protocols by Johns Hopkins University Animal Care and Use Committee (IACUC) and Department of the Army Animal Care and Use Review Office (ACURO). Rats were initially anesthetized with isoflurane (1%) via facemask and femoral intravenous and arterial catheters were inserted. Following a tracheotomy, a right-sided craniectomy was performed exposing the dura mater over the frontal-parietal cortex in all experiments with the exception of those with imaging through intact cranium. When imaging through intact cranium, the scalp was removed down to the periosteum over the right hemisphere and two burr holes (∼300 µm diameter) were created over the right somatosensory cortex spaced ∼1 mm apart for placement of a stimulating electrode pair. Following surgical preparation, the rats were placed in a stereotaxic frame and connected to a ventilator, adjusted to maintain optimal O_2_ saturation (92%-100%) and EtCO_2_ (30 – 45 mm Hg). Anesthesia continued with intraperitoneal (IP) ketamine (100 mg/kg IP) and dexmedetomidine (0.15 mg/kg IP) followed by intermittent supplementary doses as needed (25 mg/kg IP ketamine and 0.1mg/kg IP dexmedetomidine), based on physiologic changes of heart rate and blood pressure. Neuromuscular paralysis was maintained with rocuronium (7.5 mg/kg/h IP). Vital signs were monitored in real-time, and proper anesthesia and paralytic levels were maintained throughout the experiment. Standard physiological measurements including photoplethysmography (PPG), end-tidal carbon dioxide (EtCO_2_; PhysioSuite, Kent Scientific Corporation), electrocardiogram (ECG) and arterial line-recorded blood pressure (Model MP160, Biopac, Goleta, CA) at 1.25 kHz sampling rate was recorded. Temperature was monitored and maintained at roughly 37 °C via a heating pad (RightTemp, Kent Scientific Corporation). At the conclusion of experimentation, animals were sacrificed with pentobarbital (10 mg/kg IP).

### Focal Epidural Electrical Stimulation

FES (250 µA – 750 µA) was the principal methodology of neural activation. The insulated, round, single-barrel tungsten stimulating electrode pair, each with an exposed tip (<0.25 mm) were positioned (bregma: -4 mm; lateral: - 4 mm; right hemisphere) on the dural surface with approximately 1 mm separation. Care was taken to ensure electrode tips minimized dural trauma and mild tenting to maintain firm contact. A constant current isolated pulse stimulator (Model 2100, A-M Systems, Carlsborg, WA) delivered cathodic, bipolar stimulation. Typical stimulation parameters included peak current amplitudes ranging from 250 µA to 750 µA, a 5 ms pulse width, and a 10 Hz frequency for a 1 to 3 s duration.

Cortical voltage changes were monitored with an insulated, round, single barrel, stainless steel, bipolar electrode pair positioned on the parietal dural surface and recorded with EEG100C (Biopac). Timing of camera frames and event related TTL pulses were also recorded using Biopac to ensure all signals were recorded synchronously at sampling rate of 20 kHz.

### Pharmacologic experiments

General anesthesia for control experiments utilized isoflurane (0.5% – 3%). Isoflurane was administered with the ventilator via a gas vaporizer (GE Datex Ohmeda TEC 5, Chicago, IL). At least five minutes elapsed at each concentration of isoflurane, prior to presentation of the next block of FES stimuli. During control experiments with local anesthesia, topical lidocaine (4%, 50uL) was applied with a syringe to the epidural surface adjacent to the stimulating electrodes. The lidocaine was removed from the epidural surface after five minutes of exposure. For all FES control experiments, cortical EEG was monitored to track effects of anesthetic agents.

### Whisker Stimulation

For experiments imaging cortical evoked responses in wS1, epidural ECoG data was acquired simultaneous to the DHI and Biopac data using a 32-channel array (HC-32-600-10-100; NeuroNexus, Ann Arbor, MI). For these experiments, two screw electrodes were inserted on the contralateral parietal hemisphere to serve as a common reference and ground. The 3.6 mm x 3.6 mm array was composed of platinum electrodes (100 µm diameter, 600 µm spacing) on a transparent 20 µm thick polymer. ECoG data was collected at 30 kHz with a SmartBox Pro data acquisition system (NeuroNexus). Whisker deflection was achieved with a lab-built mechanical arm delivering a rostral/caudal deflection of the whisker(s) at approximately 1.5 Hz. Prior to DHI data collection, spatially distinct left-sided whiskers were identified and trimmed to a length of ∼1.5 cm with remaining whiskers trimmed to the whisker mystacial pad. Following ECoG localization in wS1 of each corresponding cortical barrel, the rodent was moved into position for simultaneous ECoG and DHI recording. Care was taken to ensure only the whisker of interest was deflected.

### Seizure

Rodent preparation was performed in a similar fashion to FES, with the only exception being the absence of stimulation electrodes. After placement of an epidural bipolar recording electrode pair (0.5 mm spacing), collection of baseline data was followed by local topical application of penicillin (Sigma-Aldrich, 6,000 u in 50 µL 0.9% saline, 1.0 mm epidural droplet application) on the dural surface. DHI was then intermittently collected to capture ictal and inter-ictal activity.

### Digital Holographic Imaging System

All optical data was collected utilizing a DHI system in an off-axis configuration designed to measure spatially resolved phase changes of a sample in reflection geometry. A laser was split between a reference arm and a sample arm. A 3 mm diameter, collimated beam illuminated the sample on-axis at skin safe operating power. Light scattered off the sample was collected and relayed through a telecentric lens system forming an image 75 mm in front of the camera. This light was then mixed with a reference beam using a beam splitter, forming a conventional off-axis hologram. The reference arm contained an optical delay line, the length of which was modified to in order to match the optical path length of the object arm with the reference arm for a targeted imaging depth. The complex image information was retrieved following the standard Fresnel image reconstruction method and the angle of the reference beam was such that the portion of the object illuminated by the 3 mm diameter beam was reconstructed in a single quadrant of the reconstructed image space, spatially separated from the autocorrelation term.

For 780 nm DHI system, holograms were acquired using a Genie Nano camera (Teledyne DALSA, Waterloo, Ontario, Canada). The camera was operated with a focal plane array (FPA) size of 256 × 256 to achieve a frame rate of 1550 Hz. The spatial resolution of the reconstructed image was approximately 30 μm, which was based on the distance from the FPA to the image formed by the telecentric lens of 50 mm and a camera pixel pitch of 4.8 μm. The 780 nm laser operated with a 3 mm coherence length, resulting in an approximately 1 mm depth resolution based on the refractive index of tissue, which was set to 1.34.

For the 1310 nm DHI system, holograms were acquired with a CRED3 camera (Axiom Optics, Somerville, MA). The camera was operated with a focal plane array (FPA) size of 256 × 256 to achieve a frame rate of 1,980 Hz. The spatial resolution of the reconstructed image was 30 μm, which was based on the distance from the FPA to the image formed by the telecentric lens of 90 mm and a camera pixel pitch of 15 μm. The 1310 nm laser operated with a 50 μm coherence length, resulting in an approximately 50 μm depth resolution.

### Data Processing and Analysis

All data processing and analysis was performed in MATLAB 2022b (MathWorks, Natick, MA).

#### Velocity Waveform Calculations

The velocity timeseries waveforms were computed with equations (1) – (3). For the FES data, the ROI was defined as a circle with a 0.5 mm diameter. Full-field maps for FES data were computed with overlapping ROIs shifted by one pixel via spatial convolution. Values outside the FOV were not included in the full-field maps. For the wS1 or seizure data, the ROI was defined as a circle encompassing the field-of-view.

For the wS1 data, the velocity waveforms were filtered with a six-pole zero-phase Butterworth bandpass filter with a cutoff frequency of 100 Hz to 500 Hz. For the through intact cranium data, velocity was bandpass filtered with a cutoff frequency of 0.5 Hz to 450 Hz.

#### Stimulus-Locked Averaging

To understand the characteristic change in tissue velocity associated with neural activity and reduce the phase noise, velocity SLA 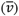 was computed by segmenting the velocity waveform into epochs based on the onset times of the stimulus. The segmented velocity (*v*_s_) was averaged across the stimulus events (*s*) at each time point, i.e.,

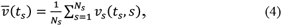

where *N*_*s*_ is the number of stimulus events and *t*_*s*_ is epoch time basis where *t*_*S*_ ∈ [*t*_*pre*_, 0] and *t*_*S*_ ∈ [0, *t*_*post*_] is the pre-stimulus baseline and post-stimulus time ranges, respectively. For wS1 data (Fig. 4D), the envelope of 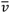 was computed as the magnitude of the Hilbert transform.

#### Pre-stimulus Z-score Normalization

Pre-stimulus Z-score normalization was applied to the segmented velocity to account for the effects of optical scattering while imaging at depth. The normalization is given by:

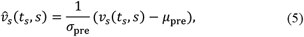

where *μ*_pre_ and *σ*_pre_ are the mean and standard deviation of *v*_*s*_ over the pre-stimulus time range, i.e., *t*_*S*_ ∈ [*t*_*pre*_, 0], respectively. For through intact cranium FES experiments, the pre- and post-stimulus time range was 10 ms. For wS1 data the pre- and post-stimulus time range was 25 ms.

#### RMS Velocity Calculations

To quantify the overall amplitude of the measured optical response, the RMS velocity over the post-stimulus time range (defined by *t*_*S*_ ∈ [0, *t*_*post*_]) was computed on the segmented velocity and with SLA, denoted by *v*_RMS_ and 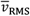 in the figures, respectively.

#### Displacement Calculations

The tissue displacement was computed by numerically integrating the velocity, given by equation (3). The total displacement was computed as the range of tissue displacement during the stimulus period.

For the seizure model, the total displacement was computed as the range of tissue displacement during an ictal discharge as manually identified by the EEG sharps.

#### Statistical Tests

Data was assessed for normality using Shapiro-Wilk test prior to hypothesis testing. For exposed cortex FES data, comparison between stimulation current conditions was performed with a Kruskal-Wallis test followed by a post-hoc Dunn–Šidák correction for multiple comparisons. For the exposed cortex FES pharmacologic data, comparisons between a given condition and the baseline condition (e.g., pre-delivery of pharmacologic) was performed with one-way ANOVA followed by a post hoc Dunn–Šidák correction multiple comparisons. For wS1 data, comparison between the pre-stimulus and post-stimulus RMS velocity was performed with a Mann-Whitney U test. For the displacement data, comparison between stimulation paradigms was performed with a Kruskal-Wallis test followed by a post-hoc Dunn– Šidák correction for multiple comparisons. The level of significance was set at p < 0.05 for all statistical tests.

#### Data analysis of electrocorticography

ECoG high gamma data processing involved computing the power of high frequency components at each electrode comparable to those frequency ranges considered for the optical data. Frequency ranges considered for optical data in lower SNR environments were determined by overcoming physiological clutter. The frequency ranges considered (80 – 350 Hz) include the “high gamma” range which has been found to be associated with neuronal activation in the cortex (36). To quantify this high frequency power over time, at each electrode in the ECoG grid the fast frequency transform (FFT) was computed using a Hamming window over 13 ms intervals slid by 0.8 ms. For each window, the average spectrum magnitude between 80 and 350 Hz was computed. These waveforms at each electrode were then averaged over those electrodes corresponding to the spatial extent of the optical response.

## Acknowledgments

We thank T. Tung and S. Cao for help in surgical planning, preparation support during rat experiments. We also thank, R. Koehler and C. Scholl for valuable scientific discussions. We thank J. Wathen and S. Rogers for their knowledge and assistance in laser support and coherent imaging; R. Hingorani and M. Fifer for knowledge and analysis of electrocorticography.

## Author Contributions

Conceptualization: DWB, MM, CLR, ATL

Methodology: DWB, MM, CLR, ATL

Software: EBK, NES

Resources: DWB, MM, CLR, ATL

Formal analysis: EBK, NES, ATL

Investigation: DWB, MM, CLR, ATL

Data Curation: EBK, NES, ATL, CLR

Project Administration: DWB, MM, CLR, ATL

Supervision: DWB, MM

Visualization: EBK, NES, ATL

